# RAVE: comprehensive open-source software for reproducible analysis and visualization of intracranial EEG data

**DOI:** 10.1101/2020.06.02.129676

**Authors:** John F. Magnotti, Zhengjia Wang, Michael S. Beauchamp

## Abstract

Direct recording of neural activity from the human brain using implanted electrodes (iEEG, intracranial electroencephalography) is a fast-growing technique in human neuroscience. While the ability to record from the human brain with high spatial and temporal resolution has advanced our understanding, it generates staggering amounts of data: a single patient can be implanted with hundreds of electrodes, each sampled thousands of times a second for hours or days. The difficulty of exploring these vast datasets is the rate-limiting step in discovery. To overcome this obstacle, we created RAVE (“R Analysis and Visualization of iEEG”). All components of RAVE, including the underlying “R” language, are free and open source. User interactions occur through a web browser, making it transparent to the user whether the back-end data storage and computation is occurring on a local machine, a lab server, or in the cloud. Without writing a single line of computer code, users can create custom analyses, apply them to data from hundreds of iEEG electrodes, and instantly visualize the results on cortical surface models. Multiple types of plots are used to display analysis results, each of which can be downloaded as publication-ready graphics with a single click. RAVE consists of nearly 50,000 lines of code designed to prioritize an interactive user experience, reliability and reproducibility.

## Introduction

The importance of high-quality software tools in advancing human neuroscience research is self-evident. The availability of purpose-built open-source fMRI software packages such as AFNI (Cox, 1996; Saad et al., 2006) and FSL (Smith et al., 2004) allowed thousands of neuroscientists and psychologists who were not experts in signal processing to analyze data from fMRI experiments. Similarly, while innovations in amplifier electronics slashed the cost of scalp encephalography (EEG) the development of comprehensive, freely-available analysis packages such as EEGLAB (Delorme and Makeig, 2004; Martinez-Cancino et al., 2020) played a key role in enabling EEG discoveries.

The last decade has seen an exponential increase in the numbers of studies investigating neuroscience questions using invasive recordings from the human brain (reviewed in Parvizi and Kastner, 2018). Recordings using grids of electrodes that sit on the cortical surface are referred to as electrocorticography (ECoG) while studies using depth electrodes that penetrate into the brain with recording contacts spaced at regular intervals along the shaft are referred to as sterotactic EEG (sEEG). Taken together, both recording techniques are referred to as intracranial EEG (iEEG). Because iEEG electrodes are implanted directly in the brain, iEEG recordings feature high spatial and temporal resolution, excellent signal-to-noise ratio and long continuous recordings without artifacts. As a result, iEEG datasets differ markedly from fMRI and EEG datasets. The development of RAVE was prompted by the limited options for neuroscientists in need of a comprehensive, open-source software package that handles all aspects of iEEG data analysis and visualization.

The design philosophy of RAVE is based on five principles. The first principle is *rigorous statistical methodology*. The history of fMRI has been marked by tumult over problematic statistics, such as “voodoo correlations” resulting from biased analyses (Simmons et al., 2007; Vul et al., 2009); treating subjects as fixed *vs*. random effects (Mumford and Nichols, 2009); and the “dead salmon” debate over how to correct for multiple comparisons (Bennett et al., 2009; Eklund et al., 2016; Cox et al., 2017). To encourage statistical best practices, RAVE is developed using “R”, a free, open source statistical language with a rich framework of existing packages developed by leading statistical and machine learning researchers (R Computing, 2017). RAVE implements robust statistical tests, such as linear mixed-effects models, in a rigorous and community-vetted fashion (Bates et al., 2015; Kuznetsova et al., 2017).

The second principle is to *keep users close to the data* so that users may make discoveries about the brain without being misled by artifacts. This necessitates a well-designed graphical-user interface (GUI) so that users can explore very large iEEG datasets combined with efficient algorithm implementation to display analysis results quickly enough for real-time, interactive interrogation.

The third principle is to *run anywhere*. RAVE is designed so that all user interactions can take place within a web browser. This makes RAVE platform and processor independent, running on tablets, desktops, or clusters. The RAVE front-end experience is the same whether data and computing resources are located on the user’s own machine, a lab server, or a cloud-based computing service such as Amazon Web Services.

The fourth principle is to prioritize *reliability and reproducibility* in all aspects of RAVE development. Funding agencies and journals have recognized that data sharing is a key ingredient in speeding scientific progress, with all research programs supported by the United States National Institutes of Health Brain Research through Advancing Innovative Neurotechnologies (BRAIN) required to submit their research data to an approved archive (Zhan, 2019). Archive submission often entails a laborious collection and curation process on the part of investigators. RAVE simplifies this process by automatically harmonizing all iEEG data files from each participant, together with meta-data such as task epoch files, in a format acceptable to archives. All processing commands used to generate a particular file or analysis can be easily documented for attachment to manuscript figures.

The final principle is to *play well with others*. Each laboratory has its own ecosystem of software tools and methods with expertise and protocols developed over years of experience. Therefore, RAVE is designed to integrate seamlessly with existing workflows, such as iELVis (Groppe et al., 2017) and `img_pipe` (Hamilton et al., 2017) for electrode localization. Along with an extensive GUI, RAVE provides a robust application programming interface (API) to support integration of RAVE with existing or novel analysis pipelines.

## Methods and Results

RAVE is freely available from the GitHub repository at https://github.com/beauchamplab/rave and currently consists of almost 50,000 lines of code and documentation. The installation also includes the option to download sample data so that users may recreate all analyses described in this manuscript. RAVE consists of interpreted R code and uses the R Shiny package (Chang et al., 2019) to create interactive web apps. The user’s interactions with these web apps via a web browser constitutes the RAVE graphical user interface (GUI). RAVE has been tested on Mac, Windows, and Linux platforms. A Docker version of RAVE is available for easy cross-platform installation of RAVE, even on computers without web access.

### Deployment Strategies

The user always interacts with RAVE using a web browser on a local machine (desktop, laptop or tablet). The location of the data repository and the hardware used for analysis computations may be independently configured, creating a variety of possible deployment strategies (Figure 1).

**Figure 1.**
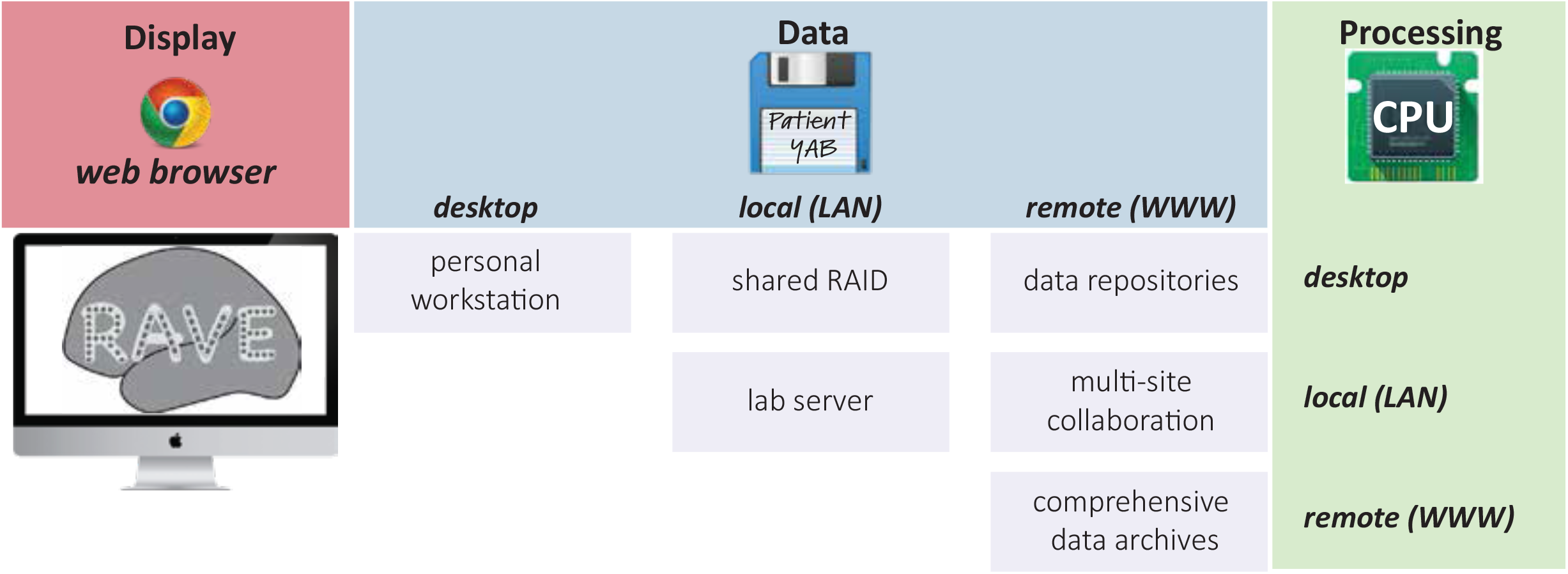
Schematic of RAVE deployment configurations. Users always interact with RAVE via a web browser running on a desktop, laptop, or tablet (“Display” column). Data storage (blue, columns) and processing (red, rows) can both occur on the same desktop machine used for display. This is mandatory in situations with little or no internet connectivity, such as a hospital room or an airline flight. In laboratory situations, processing can occur on the desktop while data storage is shared, or both processing and storage can be shared. For large-scale collaborations, data can be stored in cloud and processing can occur either on the desktop, locally, or in the cloud.

In hospital environments, internet access is usually limited (an investigator preparing a manuscript on an airplane is similarly challenged). For these situations, RAVE would run on a local machine such as desktop or laptop with the user interactions taking place within a web browser on the same machine. Both data and analysis computations would take place on the local machine.

In laboratory environments, inexpensive shared data storage is often used so that lab members to access data from many different projects and subjects (a 20 TB RAID can be purchased for less than $1000USD). In this scenario, RAVE runs on each user’s local machine, controlled via web browser. All analysis takes place locally, but data is stored on the shared RAID (typically accessed via a mount point).

A third type of deployment places analysis at a laboratory level on a shared lab compute server, such as a cluster. In this configuration, multiple RAVE sessions are run in parallel on the server, controlled by a web browser on each user’s local machine. Analysis and storage take place on the central compute server/RAID. This configuration has two advantages. First, compute servers can be equipped with dozens of cores and TB of RAM, increasing processing speed. Second, large data files are transferred exclusively over fast connections between the compute server and the RAID (both of which would often be hosted in a university data center) rather than slower desktop connections between the data center and the desktop. Network performance for desktop connections can vary, especially for telework situations in which the user is connecting via virtual private network (VPN).

Other deployments are also possible. An investigator might upload a large library of iEEG data to an online repository (Miller, 2019). Users running RAVE on their local machine could set the data location to the online repository, resulting in local analysis with remote storage. Alternately, in a multi-lab collaboration, a private data repository could hold all data collected across labs, with each lab’s compute server connecting to the repository for laboratory-level analysis with remote storage.

A final deployment strategy places both analysis and storage “in the cloud”. This would be appropriate for a comprehensive data repository, such as those under construction for the NIH BRAIN Initiative including NEMAR (Franklin, 2019) and DABI (Toga et al., 2019). A user would run a web browser on their local machine and point it to an instance of RAVE running on an academic or commercial cloud computing service, which would also host the entire data archive.

## Software Architecture

Figure 2 shows a flowchart of the components of the RAVE software most relevant to users. Using the GUI or command line calls, the user identifies the location of raw iEEG data files. For visualization, users can also identify the location of MRI images of the patient’s brain and cortical surface models created from the MRI images with FreeSurfer (Dale et al., 1999; Fischl et al., 1999) or other reconstruction tools. If a subject MRI is not available, data can be displayed on a standard template brain.

**Figure 2.**
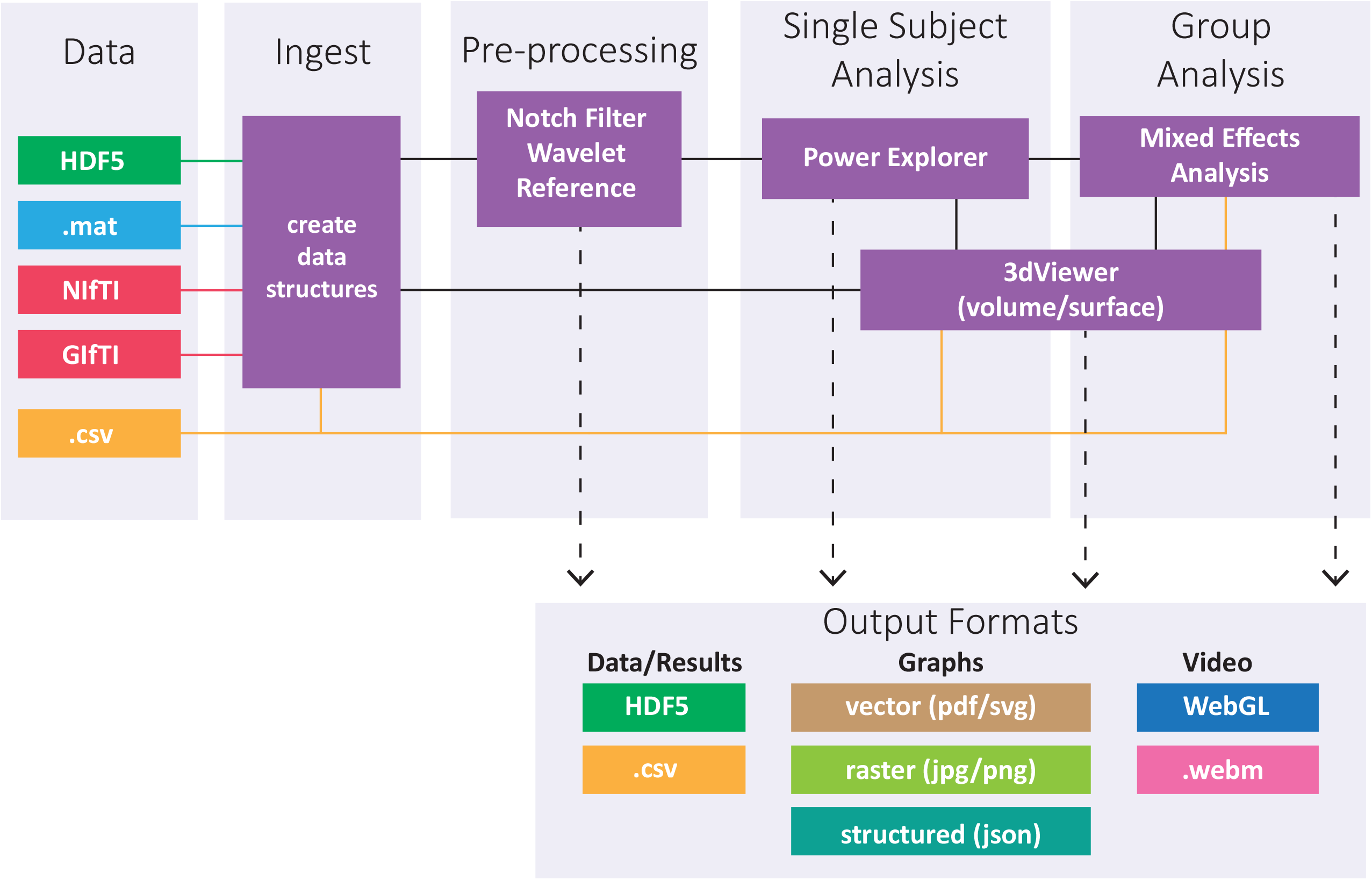
Flowchart of RAVE software architecture, proceeding from data input, through different processing stages, and ending in output of processed data and results, graphs and plots, and videos. Data files are stored in the HDF5 format for hierarchical data organization, the same format used by Neurodata Without Borders (Teeters et al., 2015) and can be imported from Matlab (.mat) format, the export format of choice for many data acquisition systems. MRI volumes are imported in standard NIfTI format and cortical surface models in GIfTI format. Comma-separated variable (.csv) files are used for human-readable tabular data, such as electrode ROI locations.

A typical iEEG patient might perform many different tasks in the course of their hospitalization. For instance, in subject 1, some data files might be a speech task, others a vision task, and still others a memory task, while in subject 2, the same tasks might be run in a different order. RAVE reorganizes all data files into a consistent structure, using a directory tree with project (such as speech, vision or memory) at the highest level, followed by subject, followed by data files. Data files are stored in the HDF5 format for hierarchical data organization, the same format used by Neurodata Without Borders (Teeters et al., 2015). As detailed below, spectral decomposition is performed by the RAVE pre-processor, followed by data exploration in single subjects with the *Power Explorer* module and across subjects with the *Group Analysis* module. In *3dViewer*, users can click on individual electrodes and view their activity profile, localizing them on the cortical surface for ECoG electrodes or on a 3-plane view of the MRI volume for sEEG electrodes.

A variety of output options are available. High-quality PDFs or PNGs of any graph or brain image can be exported with a single click from each module for use in figures within presentations, manuscripts, and grant proposals. The data underlying any analysis can be exported in text or Microsoft Excel format with a single-click to create manuscript tables or for analysis outside of RAVE. Animations showing brain activity evolving over time can also be generated easily (in *.webm* format), or even exported as a standalone web app complete with all functional and anatomical data (HTML and Javascript using WebGL).

To further the design principle of playing well with others, there are multiple entry and exit points for the processing stream: users are not locked into a sequential analysis that begins with raw iEEG data and ends with an activity plot. For instance, a clinician could visually inspect the clinical recordings from each electrode and assign an index of epileptiform activity to each electrode in a spreadsheet table. The clinician could then import this table into RAVE and display it on the cortical surface and 3-plane viewer, completely bypassing the preprocessing and analysis of the voltage-by-time data. Alternately, a data scientist could use the RAVE data structures and preprocessing modules to quickly and easily generate a single value for each condition in each trial in each electrode, then import this data into a machine learning toolbox for training and testing with leave-one-out analyses.

### Single subject analysis

The heart of the RAVE user experience is fast and interactive data exploration. While signals from neighboring scalp electrodes in EEG (or sensors in MEG) are usually similar, neighboring electrodes in iEEG often respond completely different as the electrode grid traverses functional boundaries in cortex (for ECoG) or the electrode shaft penetrates different subcortical nuclei (for sEEG). This makes accurate visualization, selection and display of individual electrodes a critical step in iEEG analysis. RAVE accomplishes this task with a dedicated module, termed *3dViewer*, for electrode visualization and selection. ECoG electrodes are displayed on a cortical surface model of the participant’s cerebral hemispheres created by FreeSurfer or another reconstruction tool (Dale et al., 1999; Fischl et al., 1999). The *3dViewer* cortical surface model view can be zoomed or rotated with the mouse or keyboard shortcuts (Figure 3A). Users can view the MRI volume in separate panels or overlaid on the cortical surface module and scroll through the slices (Figure 3B). Individual electrodes are clickable, which then updates the current analysis to focus on that electrode.

**Figure 3.**
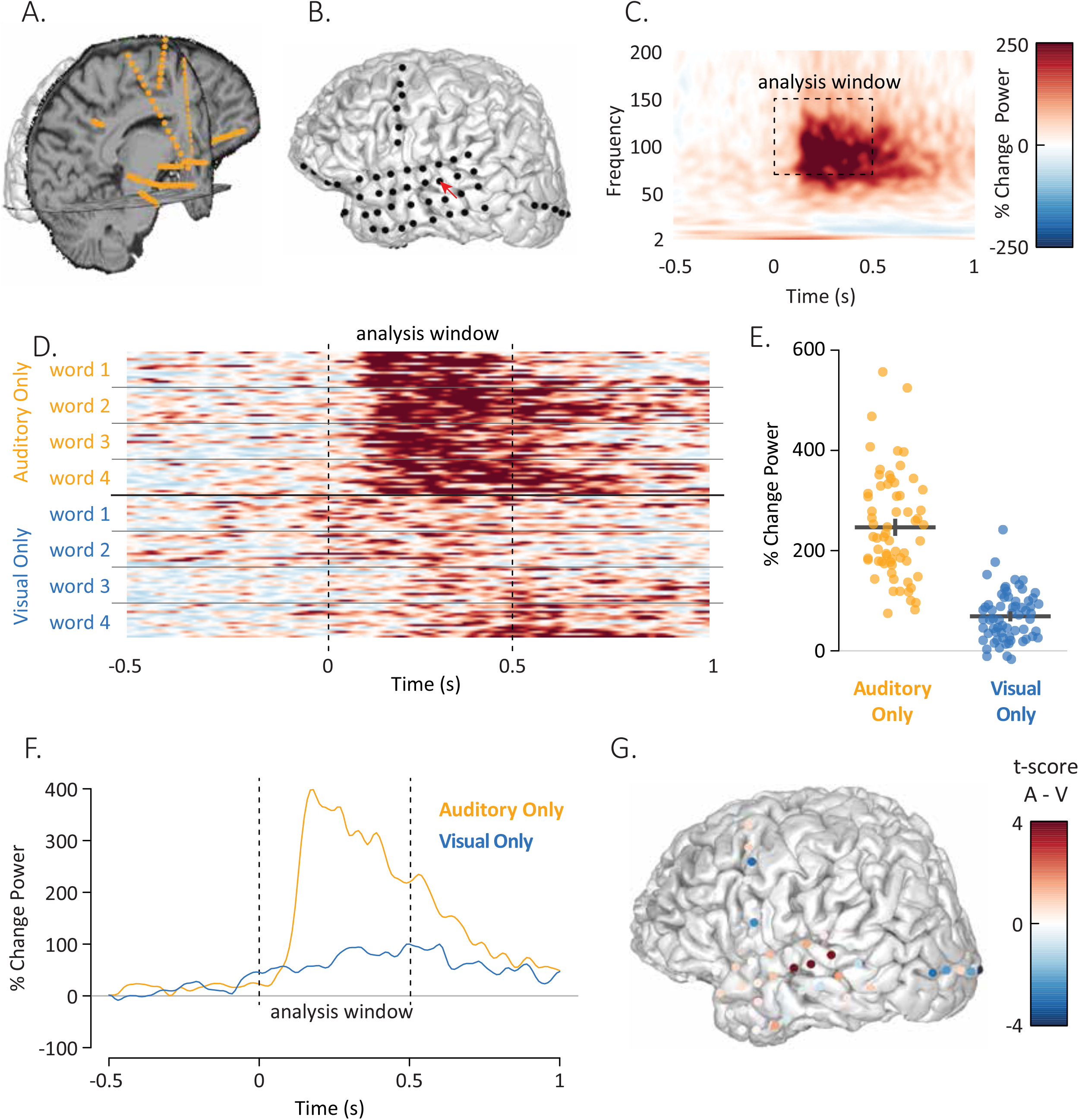
A. The *3dViewer* displays axial, sagittal and coronal slices through the MRI volume for localization of sEEG electrodes (orange spheres). B. The *3dViewer* also supports display of cortical surface models for localization of ECoG electrodes (black spheres). Red pointer highlights YAB electrode 14. C. Power-by-frequency plot showing the response to single words in YAB electrode 14. The experiment consisted of repeated presentations of single words, time zero corresponds to the onset of the auditory component of the word. D. Individual trial data from YAB electrode 14. Each row/strip shows the response in a single trial over time, collapsed across the frequencies within the analysis window. There were 8 total stimuli, 4 stimuli consisted of auditory-only recordings of words, 4 stimuli consisted of silent visual-only videos of the same words. There were 16 presentations of each individual stimulus. E. The response from each row/strip in (D) was converted to a single value by averaging over the time window from 0 seconds to 0.5 seconds. Each orange symbol shows the response to a single auditory-only trial, each blue symbol shows the response to a single visual-only trial. The black lines show the mean and standard error of the mean for each condition. F. The response to auditory-only words and visual-only words over time in YAB electrode 14, collapsed across the frequencies in the analysis window. G. The t-score of the power difference between auditory-only and visual-only conditions, calculated for each electrode and used to color each electrode.

*Power Explorer* displays several different kinds of plots to provide an in-depth view of a subject’s iEEG data. One set of statistical plots displays data for all electrodes in the subject, for a global view. The remaining plots display data for single electrodes or subsets of electrodes, selected either with the *3dViewer* or by setting anatomical or functional criteria.

The first type of plot is the time-by-frequency plot (Fig. 3C). The x-axis of the plot shows time from a given experimental event, such as the beginning of a trial or the onset of a stimulus. The y-axis shows frequency range with the power at each time-frequency cell mapped to a color scale. Both the color scale range, colors, and units of analysis (such as percent amplitude or power change from baseline, z-score of amplitude or power change from baseline, or dB from baseline) are user modifiable with a single click. For different analyses, the time-frequency range of interest differs. The range can be changed with sliders bars in the GUI, with the selected range shown as a dashed box on the spectrogram.

The iEEG signal is large enough that responses can be observed within individual trials. So that users can view the variability of the signal across individual trials, the GUI plots the response in the selected frequency range over time for each individual trial (Figure 3D). The trials are sorted by stimulus and experimental condition so that users may visually inspect the consistency of the signal and divine differences between stimuli or conditions.

The interface uses autofill text boxes so that users can group different trial types into up to 10 different conditions (the default is to group all trials into a single condition). Analysis results are instantly updated to reflect any changes. Consider an experiment with 8 trial types where each trial consists of a recording of one of four single words presented in either auditory or audiovisual format. With a few clicks, the user could group all auditory words into an “auditory” condition and all audiovisual words into an “audiovisual” condition. Alternately, each trial type could be treated as an independent condition. The composition of the conditions can be saved to disk and reloaded to avoid having to recreate complex groupings.

The activity for each defined condition is displayed in its own time-frequency plot. To display all conditions in a single plot, the data is collapsed across the time window of interest to generate a single value for each trial. Each trial is then plotted as a single point, with one column of points per condition (Figure 3E). If the user has selected more than one condition, a statistical test is automatically performed between the conditions with the results displayed above the trial plot. To compare the temporal profile of the response across conditions, the spectral signal is collapsed across the frequency range of interest and displayed, with one trace of power over time per condition (Figure 3F).

The results of the analysis across electrodes are visualized by coloring each electrode (Figure 3G). A pull-down menu permits users to select the values used for coloring from all available values for single conditions and contrasts between conditions, including raw beta-weights, t-statistics, p-values, and false-discovery-rate (FDR) corrected p-values.

### Group analysis

Discoveries made at the single subject level must be confirmed across subjects. The first step in the RAVE group analysis workflow is to identify the electrodes from each individual subject that are to be included in the group analysis, analogous to the voxel selection step in BOLD fMRI (a study of the fusiform face area might select voxels in each subject that are located in the fusiform gyrus and show a significant response to faces). In RAVE, up to three functional and two anatomical criteria can be combined to winnow down hundreds or thousands of iEEG electrodes into an appropriate subset. Anatomical labels from each electrode are taken from FreeSurfer cortical and subcortical parcellation schemes (Fischl et al., 2004) and functional criteria can be any of the statistical values generated by *Power Explorer* or other modules. For example, in a study of speech perception, electrodes in each subject might be selected using the anatomical criterion of “location on the superior temporal gyrus” and the functional criterion of “significant response to speech (p < 0.01, FDR corrected).”

An important analysis decision is the choice of the statistical threshold. To aid in this choice, RAVE displays a plot of the statistical value for each electrode along with a red dashed line showing the currently selected threshold (Figure 4A). *3dViewer* can be updated to display only the selected electrodes (Figure 4B). The entire power-by-time information for each individual trial for all selected electrodes is exported from *Power Explorer* with a single click. After this process is completed for each individual subject, the *Group Analysis* module is loaded with data from all subjects’ selected electrodes. The number and location of the electrode contributed by each subject can be visualized by coloring each electrode according to the source subject (Figure 4C).

**Figure 4.**
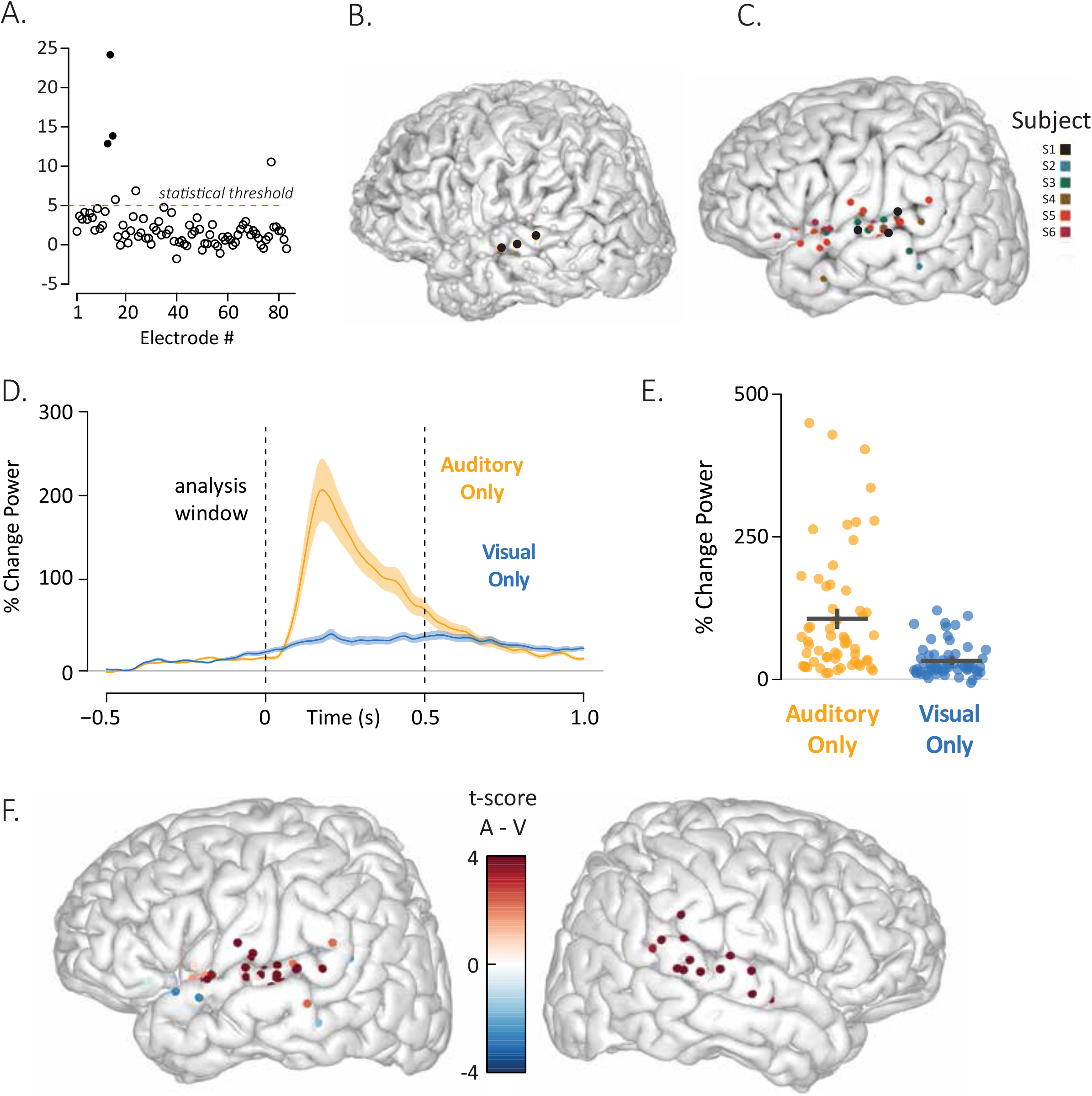
A. To visualize activity across all electrodes in a subject, RAVE plots a symbol for each electrode (x-axis is electrode number) with the y-axis displaying the result of the selected statistical test, in this case the t-test for the response to auditory-only words. Users may select electrodes for inclusion in the group analysis using up combinations of up to three functional and two anatomical criteria. The functional selection criterion is the statistical threshold (shown as a dashed red line) above which an electrode is considered responsive (t > 5 for this analysis). The anatomical selection criterion for this analysis was “located on the superior temporal gyrus”. Filled dots indicate electrodes that met both the functional and anatomical criteria, empty dots did not meet both criteria (empty dots above the red line indicate electrodes that passed the functional criterion but not the anatomical criterion). B. All electrodes in subject YAB displayed on the subject’s cortical surface model. Black electrodes met both anatomical and functional criteria in (A), gray electrodes did not. C. The same anatomical and functional criteria developed in (A) and (B) were applied to 832 electrodes in eight subjects. A total of 60 electrodes met both criteria. The locations of all left hemisphere electrodes that met the criteria are plotted on a template brain, colored by subject number (two subjects had only right hemisphere electrodes). D. The mean response across the 60 selected electrodes to auditory-only and visual-only words are plotted. Shaded regions indicate standard error across electrodes. E. The response over time in each electrode was converted to a single value by averaging over the time window from 0 seconds to 0.5 seconds. Each orange symbol shows the response of a single electrode across all auditory-only trials, each blue symbol shows the response of a single electrode across all visual-only trials. There was a significant difference between the conditions (p < 10^-16^) taken from the table showing the results of the linear mixed-effect model across all electrode and subjects. F. Results of the statistical contrast between the auditory-only and visual-only conditions was calculated for each electrode and used to color the electrodes, displayed on the left and right hemispheres of a template brain.

Because all information for all selected electrodes is loaded, the *Group Analysis* module lets users explore different analysis windows and condition contrasts for all electrodes in a study, in the same way as is possible for individual subjects in *Power Explorer.* Group contrasts and analysis windows are automatically prepopulated based on the single subject analysis settings but can be changed.

Next, a linear mixed effect (LME) model is used to perform statistical tests across all electrodes and subjects. Users can easily select any variable to serve as the dependent variable or as random or fixed effects. The default for the dependent variable is the unit of analysis (*e.g.* power change from baseline). The model defaults to treating electrode and subject as random effects, with electrode nested within subject in a multi-level fashion. Additional random effects, such as stimulus exemplar (Westfall et al., 2016) are easily added. The formula provided to the LME package is displayed, *e.g.* for the default model, y ~ 1 + (1|Subject/Electrode).

The results of the group analysis are shown in a variety of tables and plots. The complete tables includes random effects; estimates, standard errors, degrees of freedom and hypothesis test results for fixed effects in the form of an LME regression table; the omnibus fixed effects test (equivalent to the main effect of condition); the comparison of all fixed effects against zero; and all pairwise comparisons between fixed effects.

There is also a sortable and searchable table showing the univariate statistical tests for each individual electrode included in the group analysis. Rows and columns from this table can be selected in the GUI and plotted. This is important for quick examinations of response differences between regions, subjects, and any other experimental variable.

The plots available in *Group Analysis* are similar to those available in *Power Explorer*, except they are created at the group level. The power-by-time plot shows one trace per condition, but estimates of variance are calculated across all electrodes included in the analysis to provide a graphical representation of the confidence intervals (Figure 4D). To examine the consistency of the effect across electrodes, the value in each condition for each electrode is plotted (Figure 4E). An anatomical representation of the analysis results is created by coloring each electrodes by the results of the selected statistical test (Figure 4F).

A unique aspect of the *Group Analysis* module is the ability to create custom graphs. Using the “post-hoc plot” panel, the user can select “x” and “y” plot variables from a drop-down menu that contains all variables from the analysis output. A user might wish to assess whether an electrode’s response in condition 1 predicts the response in condition 2. Selecting the appropriate variables in the drop-down menu, the module creates a plot and performs the selected statistical test, including Pearson and Spearman correlations and t-test or Wilcoxon test of differences.

To assess the relationship between two variables while holding a third variable constant (partial regression), users can also select a third (“z”) variable. In an experiment with three conditions, a user might wish to assess whether an electrode’s response in condition 1 predicts the response in condition 2, partialling out the response in condition 3.

The panel also support creating variables from user entered “R” code; for example, to assess whether the *difference* in response between two conditions predicts the response in a third condition.

All plots and tables can be easily downloaded as high-resolution PDFs for manuscript or presentation figures, or as .csv files for use in manuscript tables or additional analyses.

### Preprocessing and Referencing

Because preprocessing of iEEG and EEG data are similar, the preprocessing workflow in RAVE is based on that in the widely-used EEGLAB package (Delorme and Makeig, 2004). To assess data quality, a variety of plots for each block of data and each channel are generated. RAVE creates a PDF showing data quality analytics, with one electrode per page, so that users may scan for problematic data. Analytics include plots of voltage *vs.* time after notch filtering; periodograms before and after notch filtering (both normal and log10 frequency); and histograms signal voltage. Spectral analysis is performed using wavelets with default parameters of 16 kernels with lengths from 0.101 to 1.433 seconds and cycles from 3 to 16 {Cohen, 2014 #6330}. After waveletting, data is downsampled to 100 Hz to reduce disk space; this value is also customizable. RAVE provides tools for semi-automated trial epoching based on signals present in the iEEG data, such as acquisition system analog inputs from microphones signaling the onset of auditory events or from photodiodes signaling the onset of visual events. Users can also provide .csv files that contains the times of events within each trial, such as auditory onset, visual onset, or motor response. In *Power Explorer*, a drop-down menu lists all available trial reference timepoints and analyses are time-locked to the selected event.

A key step in EEG analysis is the choice of voltage reference. RAVE incorporates three popular reference schemes: common average reference, white matter reference, and bipolar referencing. Each of these schemes has advantages and disadvantages. With the common average reference, the average of the signal at all electrodes is subtracted from the signal at each individual electrode at every time point. In some clinical situations, there may be a preponderance of electrodes over a single brain area, such as cortex important for speech or motor functions. This concentration is strongly weighted in the common average, with the result that subtracting the common average removes neural signals of interest. In practice, experimenters are well-advised to analyze their data using different schemes to understand their influence on the results.

To make this process easier, RAVE flips the usual order of operations. In most EEG processing pipelines, data is referenced and then spectral analysis is performed. However, spectral analysis is by far the most time-consuming step of the analysis and expands data volume many-fold, making it tedious and inefficient to repeatedly select a different reference and re-run spectral analysis. Instead, RAVE performs spectral analysis first, followed by referencing. Since referencing is a purely arithmetic operation, it can be performed very quickly so that users can quickly assess the effects of different reference schemes. Mathematically, both referencing and waveletting are linear convolutions, with the result that the order of operations does not change the final result as long as both real and complex components of the spectral signal are preserved.

## Discussion

RAVE provides an easy-to-use software tool so that users with no programming or signal processing expertise can analyze iEEG data and create publication-ready figures and plots.

Direct recording of neural activity from the human brain using implanted electrodes is one of the fastest-growing techniques in neuroscience. Translating the vast quantity of data collected with iEEG into neuroscience discoveries is difficult. RAVE eases discovery by providing a powerful, comprehensive, free, user-friendly toolkit that makes it easy to analyze and view iEEG data. For most existing solutions, users interact with their data by coding, often in Matlab. RAVE eliminates the necessity for programming expertise. Simply clicking on an electrode prompts immediate display of a number of useful analyses including the spectrogram, the response amplitude over time, the response to each individual trial, and the time series of the response at every trial. In contract to current solutions, in RAVE modifying any analysis parameter (such as the time-frequency analysis window) instantly updates all results. Development of RAVE is guided by five design principles.

### Rigorous statistical methodology

The linear mixed-effects model used in the *Group Analysis* module incorporates our current understanding of the most rigorous way to analyze iEEG data. Individual electrodes and subjects are modelled as hierarchical random effects. Many iEEG analyses treat electrodes and subjects as fixed effects, creating the illusion of immense statistical power since there thousands of individual experimental trials. However, the high variability across electrodes and subjects makes this analysis a poor fit to iEEG data.

No software with flexible analyses can prevent users from making statistical errors such as circular or biased analyses. However, by making the analysis transparent and easy to reproduce, other investigators to easily examine claims based on iEEG data and determine if the underlying methodology is sound. Furthermore, for users who rely on the RAVE GUI, all of the code is already shared. This permits community-based examination of the internal workings of RAVE and for any errors that are discovered to be corrected. This is sharp contrast to current practice, where each trainee in a single laboratory might analyze data using a different combination of functions called in different order with different parameters. This can lead to inconsistency if not outright errors and make it very difficult to reproduce analyses. *Keep users close to the data*

The *Power Explorer* module is designed for interactive data visualization to power new discoveries. Data is automatically displayed broken down by individual trials, sorted by conditions. This is important because many iEEG discoveries (as in the rest of science) are serendipitous. For instance, in an experiment originally designed to examine differences between auditory and audiovisual words, RAVE users noticed differences between different stimulus exemplars explained by the relative timing of the auditory and visual speech (Karas et al., 2019). Experimental conditions can be defined on-the-fly, with the results instantly viewable. This is more flexible than other workflows (such as generalized linear model specification in analysis of BOLD fMRI data) in which the trials making up each condition, and the contrasts between the conditions must be prespecified.

### Run anywhere

RAVE has been successfully used on Mac, Windows PCs, Linux boxes and iPads. The use of a web browser for user interfacing means that RAVE development is “future proofed” against obsolescence of specific graphics libraries, operating systems, and processing architectures.

As biomedicine moves towards funding-agency and journal enforced data archiving and sharing, the ability of RAVE to operate in the cloud will become increasingly important. RAVE automatically organized all data and meta-data for each participant into a project directory; the entire project directory can then be uploaded to a data archive, as mandated by many journals and funding agencies. The organized directory structure and analysis files created by RAVE are also easily translatable into nascent formats such as Neurodata Without Borders (Teeters et al., 2015) and the Brain Imaging Data Structure for iEEG (Holdgraf et al., 2019).

### Reliability and reproducibility

Replicating iEEG analyses can be challenging. Even if the code used for an analysis is available, changes in hardware, operating systems, and dependencies between different software tools can make it impossible to load data or execute the analysis code. A good solution to this problem is the use of the *Docker* system. “Docker distributes a binary image in which all of the software has already been installed, configures and tested, and can even include all data and metadata necessary for replication (Boettiger, 2015). can also be used for RAVE installation on hardware with limited or no internet access, frequent in patient care environments. The RAVE source code repository on github has a Docker image of a complete RAVE install.

### Play well with others

An immense amount of resources have been devoted to the development of EEG and MEG tools, such as EEGLAB (Delorme and Makeig, 2004), FieldTrip (Oostenveld et al., 2011) and MNE (Gramfort et al., 2013; Gramfort et al., 2014). RAVE is designed to integrate with these tools via data exchange through open file formats or direct function calls via API for access to internal data structures. Together with `Rcpp`, an R interface to the programming language C++, `reticulate`, an R interface to Python, and the `R.matlab` package, an interface to Matlab, the API permits RAVE users to create their own analysis modules in a variety of languages.

All components of RAVE import or export comma-separate value (.csv), making data exchange simple. For example, there are a number of processing pipelines for localizing iEEG electrodes (RAVE also includes rudimentary tools for this purpose). More sophisticated solutions, such as iELVis (Groppe et al., 2017) and `img_pipe` (Hamilton et al., 2017) generate .csv files with one row per electrode. RAVE directly reads these files and uses them to locate electrodes. On the export side, *Power Explorer* generates .csv files, allowing users to use RAVE for initial stages of analysis and then export the data for more complex analyses not implemented in RAVE, such as machine learning.

## Acknowledgments

This research was supported by NIH R24MH117529.

We thank Meng Li for statistical advice and are grateful for feedback from RAVE users including Anusha Allawala, Kelly Bijanki, Patrick Karas, Brian Metzger, Buffy Nesbitt and Sameer Sheth.

